# ARHGAP10 is a novel microtubule-associated protein that regulates the resorption activity of osteoclasts

**DOI:** 10.1101/2025.06.12.659268

**Authors:** Laura Jentschel, Anne Blangy, Guillaume Bompard

## Abstract

Adult-bone homeostasis is maintained through the reciprocal actions of osteoclasts and osteoblasts, which respectively resorb and deposit new bone. Excessive osteoclast activity leads to bone loss and contributes to conditions like osteoporosis. Osteoclasts form a specialized adhesion structure called the actin ring that is crucial for bone resorption and relies on both the actin and microtubule cytoskeletons. Our previous studies identified the β-tubulin isotype TUBB6 as a regulator of actin ring dynamics essential for osteoclast function, and found ARHGAP10, a negative regulator of the GTPases CDC42 and RHOA, as a potential mediator of TUBB6 function. Here we show that ARHGAP10 is a novel microtubule-associated protein critical for osteoclast function. ARHGAP10 directly binds microtubules through its BAR-PH domain, which requires positively-charged lysine residues K37, K41 and K44 within the BAR domain. CRISPR/Cas9 mediated knockout of *Arhgap10* affects the morphology of the actin ring and impairs osteoclast resorption activity, correlated with altered actin ring dynamics. Complementation experiments reveal that the ability of ARHGAP10 to bind microtubules and to negatively regulate RHO-GTPases are essential for its role in osteoclast resorption activity. These findings uncover a novel cytoskeletal regulator in osteoclasts and suggest that targeting the microtubule-actin interface via ARHGAP10 could represent a therapeutic strategy in bone loss disorder.

## INTRODUCTION

Adult-bone homeostasis relies on osteoclast and osteoblast activities. Osteoclasts are giant multinucleated cells, deriving from the differentiation of bone marrow macrophages, involved in bone resorption, while osteoblasts are required for new bone formation. The equilibrated balance between the activities of the two cell types is crucial for bone renewal. Indeed, excessive activity of osteoclasts can lead to progressive bone loss and potentially induces osteoporosis, as observed in several physio-pathological situations such as age-related sexual hormone decay after menopause, chronic inflammation and cancer (Khosla and Hofbauer, 2017). Upon polarization, osteoclasts organize at the apical face an adhesive structure in contact with the bone called the sealing zone, also known as the actin ring, composed of actin-rich, αVβ3 integrin-containing adhesive structure called podosomes (Blangy et al., 2020). The actin ring segregates a distinct membrane structure called the ruffle border, which is the site of acid and protease secretion, respectively required to dissolve underlying bone mineral and proteins. The actin ring is a dynamic structure and enables osteoclasts to slide across the bone surface and generate new resorption pits (Søe and Delaissé, 2017).

While many efforts have focused on inhibiting osteoclast differentiation as a strategy to prevent excessive bone degradation, we have recently demonstrated that targeting the actin ring represents a relevant therapeutic approach against osteoporosis (Blangy et al., 2020; Mounier et al., 2020; Vives et al., 2015). Therefore, it is critical to understand the mechanisms regulating actin ring formation and dynamics, in order to identify potential new targets. To address this question, we performed a differential transcriptomic / proteomic study that led to the identification of candidates among which TUBB6, an isotype of β-tubulin (Guérit et al., 2020). Actin ring formation and dynamics do not rely solely on the actin cytoskeleton but also on microtubules, which regulate podosome organization (Destaing et al., 2003; Morel et al., 2024). Thus, the two cytoskeletons are intimately associated to promote bone resorption by osteoclasts. We investigated the role of TUBB6 and demonstrated that it plays a key role in osteoclast activity (Maurin et al., 2021). In absence of TUBB6, osteoclasts display actin ring defects correlated with changes in microtubule but also actin dynamics resulting in inhibition of resorption activity (Maurin et al., 2021). In order to identify proteins potentially linking TUBB6 to actin cytoskeleton dynamics, we performed a differential proteomic analysis of fractions enriched in microtubule-associated proteins (MAPs) prepared from osteoclasts expressing or not TUBB6. From this we characterized ARHGAP10 as enriched in the microtubule fraction prepared from *Tubb6* KO osteoclasts. Furthermore, we found that ARHGAP10 colocalizes with actin and microtubule cytoskeletons in osteoclasts (Maurin et al., 2021). ARHGAP10 is a GTPase-activating protein (GAP) that negatively regulates the activity of the small GTPases CDC42 and RHOA (Ren et al., 2001; Koeppel et al., 2004); both GTPase being crucial for actin ring dynamics and osteoclasts resorption activity (Destaing et al., 2005; Ito et al., 2010). Furthermore, ARHGAP10 depletion has been shown to induce actin ring defects in osteoclasts (Steenblock et al., 2014). Taken together, these data suggest that ARHGAP10 may be a key player in osteoclast function by mediating TUBB6 activity.

In this study, we mechanistically dissect the role of ARHGAP10 in osteoclast activity. We demonstrated that ARHGAP10 co-sediments with microtubules via its amino-terminal BAR-PH domain. We identified critical residues involved in the direct binding of the BAR-PH domain to microtubules by modeling it. The role of ARHGAP10 in osteoclast activity was studied by knocking-out *Arhgap10* in RAW264.7 cells. RAW464.7 cells are osteoclast precursors: when grown in the presence of the cytokine RANKL, they differentiate into multinucleated bone resorbing cells with the characteristics expected of fully differentiated osteoclasts (Collin-Osdoby and Osdoby, 2012). We used crystalline inorganic calcium phosphate dissolution assays to assess osteoclast activity. In fact, extracellular medium acidification, which leads to crystalline inorganic calcium phosphate dissolution, is a typical property of osteoclasts that relies on an intact actin ring. It is the first step of the bone resorption process and it is necessary to make bone proteins, which are embedded in hydroxyapatite, amenable for degradation by proteases (Blangy et al., 2020). Using this model, we showed that actin ring organization and dynamics were affected upon depletion of ARHGAP10, as was osteoclast activity. Finally, we demonstrated that microtubule-binding capacity, as well as GAP activity, of ARHGAP10 are required for osteoclast function.

## RESULTS

### ARHGAP10 is a new microtubule-associated protein

We previously found that upon osteoclast fractionation, the endogenous ARHGAP10 protein was present in a fraction containing microtubule-associated proteins (MAPs). Furthermore, ARHGAP10 colocalizes with the actin ring and with microtubules in osteoclasts, and that it also co-sedimentes with microtubules when overexpressed in HEK293T cells (Maurin et al., 2021). In order to test whether ARHGAP10 could be a genuine MAP, we first performed a microtubule co-sedimentation experiment using high speed supernatant from osteoclasts whole cell extracts. After the induction of tubulin polymerization, microtubules and associated proteins were sedimented. Immunoblot analysis demonstrated that endogenous ARHGAP10 equally fractionated in the supernatant and in the pellet containing microtubule (Fig. 1A). The efficacy of the fractionation was assessed by the presence of the non-MAP protein vinculin only in the supernatant and whereas GEFH1 exclusively co-sedimented with microtubules as expected from its known capacity to bind microtubules, including in osteoclasts (Morel et al., 2024). Thus, endogenous ARHGAP10 is able to co-sediment with microtubules in osteoclast extracts, suggesting that ARHGAP10 can bind microtubules.

**Figure 1:**
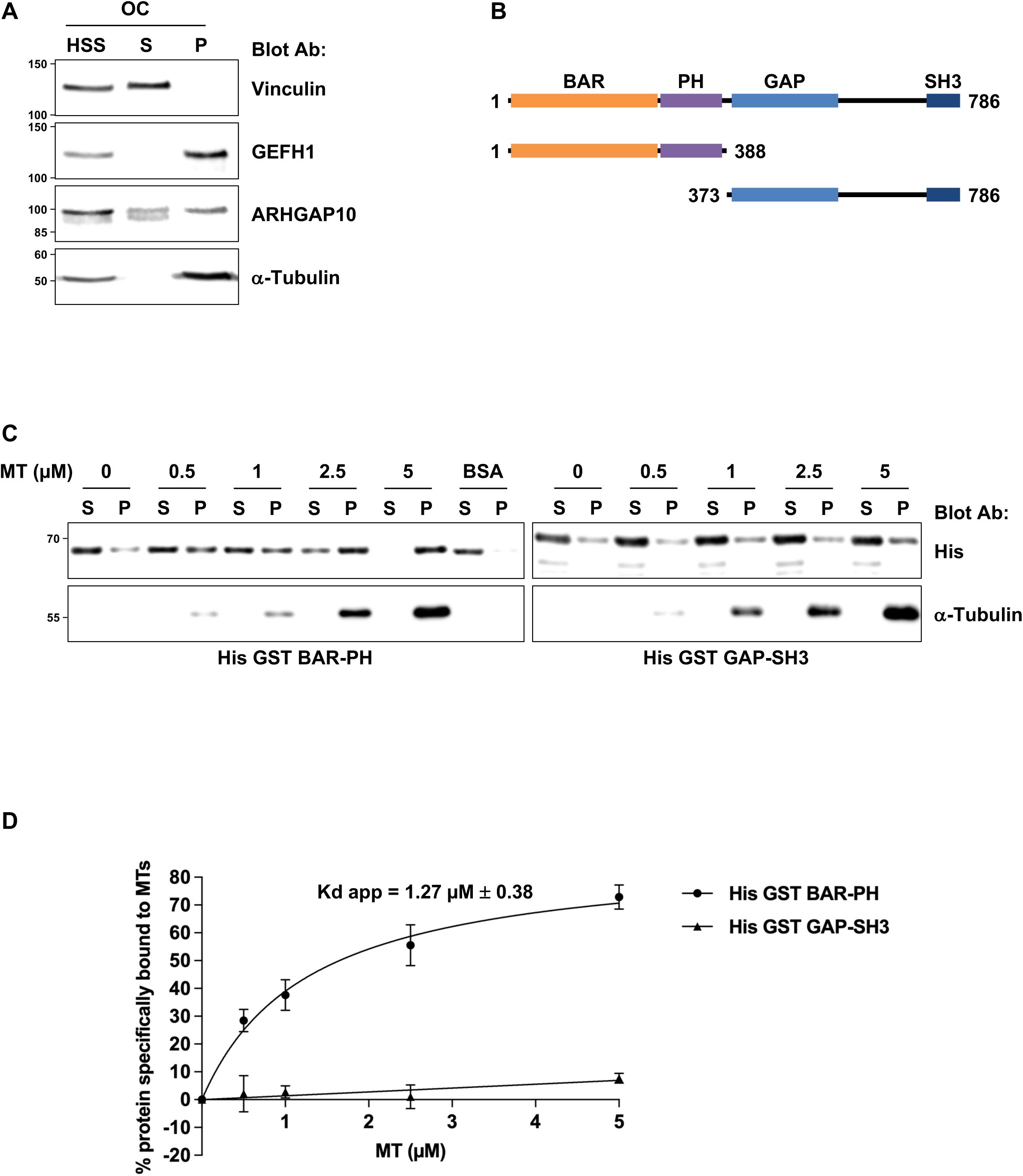
ARHGAP10 is a microtubule-associated protein. **(A)** Representative immunoblot analyses of microtubule co-sedimentation assay using high speed supernatant (HSS) from osteoclasts (OCs). A large fraction of ARHGAP10 co-sedimented with MTs (P) as GEFH1, a well characterized MAP. Vinculin is retrieved exclusively in the supernatant (S) as expected. **(B)** Schematic representation of ARHGAP10 protein with main domains depicted (BAR, PH, GAP and SH3), amino-acid numbering and fragments used for microtubule co-sedimentation assay fused to His GST. **(C)** Representative immunoblot analysis of *in vitro* microtubule co-sedimentation assay using either recombinant (1 µM) BAR-PH or GAP-SH3 fused to His GST and microtubules prepared by polymerizing pig brain tubulins. Note the increase co-sedimentation of BAR-PH with MTs in a concentration dependent manner and not with BSA demonstrating a direct binding between the two proteins. **(D)** Graph quantifying the microtubule co-sedimentation of His GST BAR-PH (n = 7, ± SEM) and GAP-SH3 (n = 3, ± SEM). The line represents the theoretical curve for a bimolecular interaction. The apparent dissociation constant (Kd app) was calculated using Prism 10.4.1 software (one site – specific binding).

To determine whether the binding of ARHGAP10 to microtubules is direct or not, we performed *in vitro* microtubule co-sedimentation experiments using pig brain tubulin and pure recombinant proteins. ARHGAP10 is composed by a BAR (*Bin, Amphiphysin, Rvs*), a PH (*Pleckstrin Homology*), a GAP (*GTPases Activating Protein*) and an SH3 (*Src Homology 3*) domains (Fig. 1B). As the full-length recombinant ARHGAP10 protein produced either in bacteria or insect cells was insoluble (data not shown), we assessed the microtubule binding properties of the N-terminal BAR-PH (aa, 1-388) and the C-terminal GAP-SH3 (aa, 373-786) regions (Fig. 1B). The BAR-PH region of ARHGAP10 co-sediments with microtubules in a concentration-dependent manner (Fig. 1C), while no co-sedimentation of ARHGAP10 was observed when BSA was used instead of microtubules. The apparent binding affinity of ARHGAP10 BAR-PH to microtubules was 1.27 µM ± 0.38 (Fig. 1D), a value in agreement with the affinity of most MAPs for microtubules (Cheeseman et al., 2006). Conversely, the GAP-SH3 domain of ARHGAP10 did not co-sediment with MT (Fig. 1C). These data demonstrate that ARHGAP10 is a new genuine MAP, which binds to microtubules through its BAR-PH domain.

### ARHGAP10 binding to microtubules requires lysine 37, 41 and 44 from its BAR domain

MAP binding to microtubules generally involves positively charged residues, as the microtubule lattice is negatively charged because of the exposition of the acidic carboxy-terminal tails of α and β-tubulins (Nogales et al., 1999; Okada and Hirokawa, 2000). In order to identify the exposed residues within the BAR-PH sequence that are positively charged, we modeled this region using SWISS-MODEL (https://swissmodel.expasy.org). The best homology hit was found when the structure of the BAR-PH of APPL1 (adaptor protein, phosphotyrosine interacting with PH domain and leucine zipper 1) was used as template (Li et al., 2007; Zhu et al., 2007). In the proposed model, two ARHGAP10 BAR-PH domains would form an antiparallel homodimer though their BAR domains, with the PH domains located at both extremities (Fig. 2A). The structure of the homodimer presents a crescent-shape characteristic of BAR domains that is required to sense membrane curvature and to induce membrane tubulation (Fig. 2A, side view) (Salzer et al., 2017).

**Figure 2:**
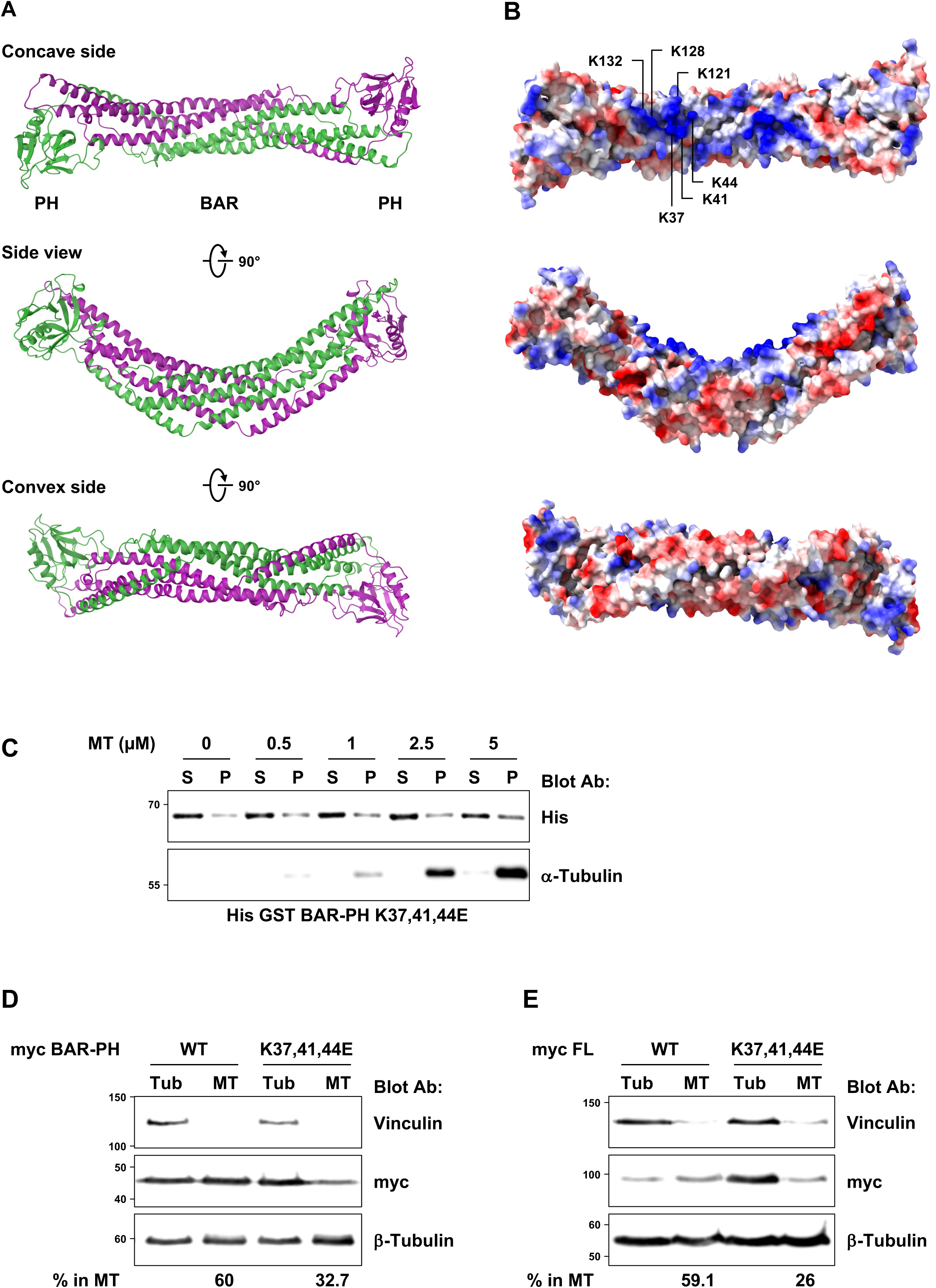
Lysine 37, 41 and 44 of ARHGAP10 are involved in MT binding. **(A)** Structural model of ARHGAP10’s BAR-PH domain. The BAR-PH domain (1-388) of ARHGAP10 was modelized using SWISS-MODEL which used APPL1 (SMTL ID: 2q13.1) as template. The BAR-PH domain of ARHGAP10 forms a homodimer, each monomer is represented in different color (green and purple). **(B)** Electrostatic surface of the proposed model. Positively charged exposed residues (blue) clustered within the concave side of the BAR-PH are indicated for one monomer. **(C)** Representative immunoblot analysis of *in vitro* microtubule co-sedimentation assay using recombinant (1 µM) BAR-PH K37,41,44E mutant fused to His GST and MT as described in figure 1C. The amount of recombinant protein co-sedimented with microtubules is not significantly affected by their concentration. Representative immunoblot analysis of lysates from HEK293T expressing myc-tagged BAR-PH **(D)** or full-length ARHGAP10 **(E)** WT or K37,41,44E subjected to tubulin/microtubule fractionation. The percentage of ectopic proteins associated with the microtubule fraction is indicated below. Vinculin is used as a marker of the tubulin fraction (Tub) and β-tubulin shows the proportion of unpolymerized (Tub) / polymerized tubulin (MT).

The electrostatic surface of the BAR-PH model for ARHGAP10 (Fig. 2B) reveals several regions of exposed positively charged residues, among which lysine residues K37, K41 and K44 as well as lysine residues K121, K128 and K132, all located within the concave side. We then evaluated *in silico* the consequences of mutating the above positively charged lysine residues to negatively charged glutamic acids (K37,41,44E and K121,128,132E) on the structure of the BAR-PH, using again SWISS-MODEL and APPL1 as a template (Fig. S1A, B). Neither the K37,41,44E nor the K37,41,44E and K121,128,132E mutations caused major changes on the overall structure of the BAR-PH homodimer, while the charge reversion was obvious on electrostatic surface models (Fig. S1A, B).

We then examined the microtubule binding capacity of ARHGAP10 BAR-PH region bearing the K37,41,44E mutations, by *in vitro* MT co-sedimentation assay. Unlike its WT counterpart, the K37,41,44E mutant was not able to bind to microtubules (Fig. 2C and Supplemental Fig. 1C). Combining the K121,128,132E and the K37,41,44E mutations did not affect microtubule binding any further (data not shown).

We then examined the consequences of the K37,41,44E mutation on the ability of the BAR-PH and full-length ARHGAP10 to bind microtubules in cells. For this, tubulin and microtubules were fractionated from HEK293T cells overexpressing myc-tagged BAR-PH or full-length ARHGAP10, WT or bearing the K37,41,44E mutations. More than half of the WT BAR-PH or full-length ARHGAP10 ectopic proteins associated with the microtubule fraction (Fig. 2D-E), similar to the endogenous protein in osteoclasts (Fig. 1A). In contrast, the K37,41,44E mutation strongly reduced the ability of both ectopic, full length and BAR-PH, ARHGAP10 proteins to co-sediment with microtubules (Fig. 2D-E).

Taken together, our data demonstrate that ARHGAP10 directly binds microtubules, involving lysine residues 37, 41 and 44 located into its BAR domain.

### ARHGAP10 regulates acting ring organization and is required for osteoclast activity

In order to evaluate the role of ARHGAP10 in osteoclast activity, we generated the knockout of *Arhgap10* in RAW264.7 cells using the CRISPR/Cas9 system. Two individual clones were isolated and characterized by immunoblot analysis (Fig. 3A), using a home-made antibody directed against the SH3 sequence of human ARHGAP10 as described (Lucken-Ardjomande Häsler et al., 2020). Neither *Arhgap10* KO clone expressed detectable ARHGAP10 (Fig. 3A) but both retained the ability to differentiate into osteoclasts, as assessed by TRAP staining (Fig. 3B). We then examined the consequence of ARHGAP10 ablation on actin ring organization. For this, osteoclasts were cultured on mineral matrix and the structure of the actin ring was classified according to categories, according to their aspect: normal, abnormal, multiple or absent (Fig. 3C), as described the material and methods section. The actin ring was normal in around 60% of WT osteoclasts, whereas this value dropped by half in osteoclasts devoid in ARHGAP10 (Fig. 3D). This was correlated with an increase in the proportion of osteoclasts presenting actin ring defects. More *Arhgap10* KO osteoclasts presented multiple small actin rings or an abnormal fragmented sealing zone at the expense of normal ones, while the percentage of osteoclasts without actin ring was barely affected (Fig. 3D). Such actin ring defects are usually associated with osteoclast resorption defects. In aggreement with this, we observed that *Arhgap10* KO osteoclasts showed a severe reduction on their capacity to dissolve hydroxyapatite (Fig. 3E).

**Figure 3:**
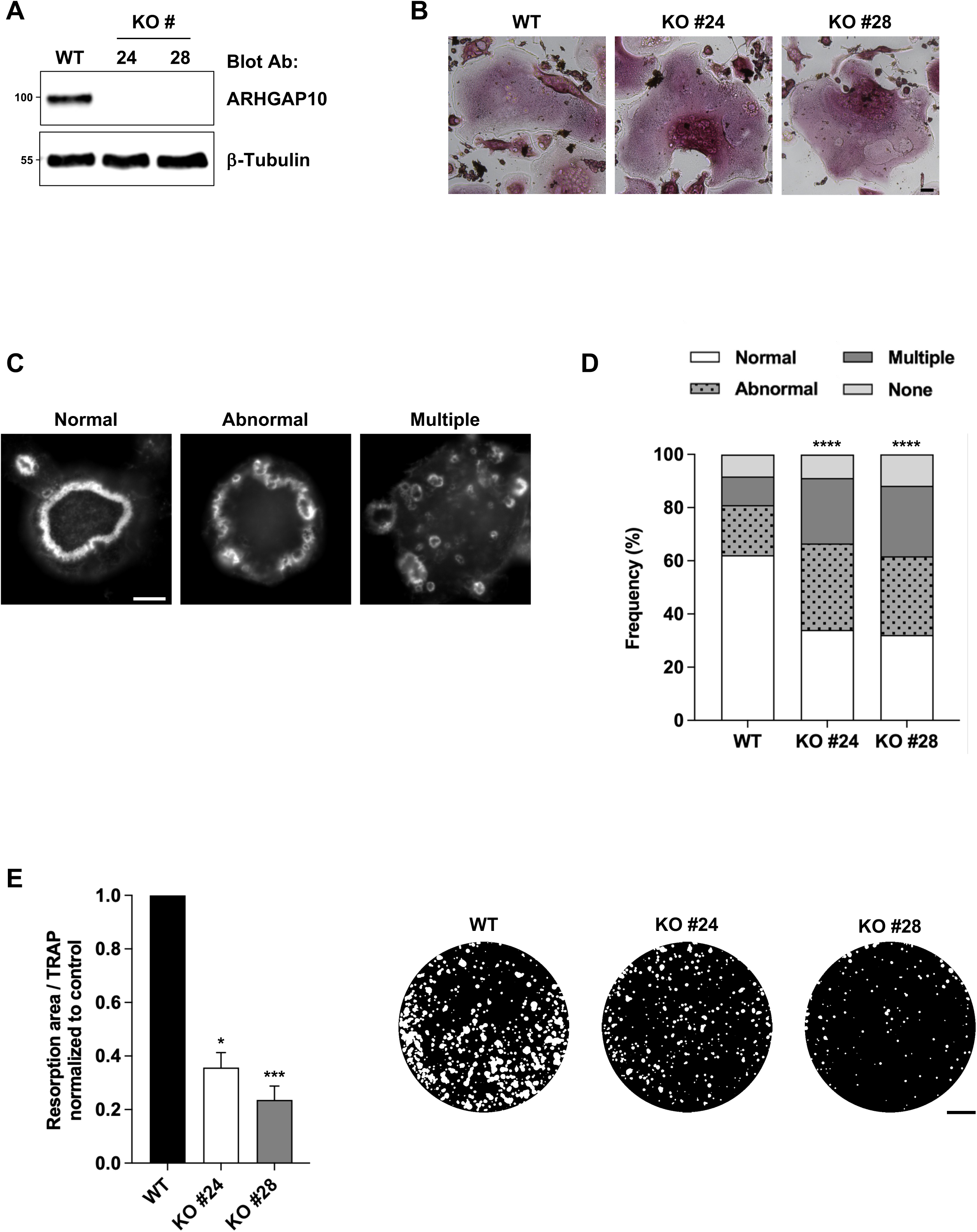
ARHGAP10 regulates osteoclast activity. **(A)** Representative immunoblot analysis of RAW264.7 cells KO for Arhgap10. β-tubulin shows equal protein loading. **(B)** Representative images showing TRAP staining in WT and *Arhgap10* KO osteoclasts. Bar, 50 µm. **(C)** Representative images illustrating the different categories of actin ring organization considered in (D). Bar, 10 µm. **(D)** Bar graph showing the frequency of WT and *Arhgap10* KO #24 or 28 osteoclast seeded on mineral matrix and presenting normal, abnormal, multiple, and absent (none) actin ring. A minimum of 300 osteoclasts per genotype from three independent experiments were counted, χ^2^ contingency test, ****p < 0.0001. **(E)** Graph showing the average specific resorption activity of Arhgap10 KO osteoclasts normalized to the activity of WT osteoclasts (n = 10, ± SEM; 8 wells analyzed per experiments), Friedman test: *p < 0.05 and ***p < 0.001. Representative images of von Kossa staining of the resorption wells are presented on the right. Bar, 1 mm.

### Consequences of ARHGAP10 on actin dynamics in osteoclasts

The actin ring and resorption defects of *Arhgap10* KO osteoclasts were reminiscent of those we reported formerly upon *Tubb6* KO, in which actin dynamics was also altered (Maurin et al., 2021). Thus, we assessed the impact of *Arhgap10* KO on the dynamics of actin rings, by performing time lapse videomicroscopy in osteoclasts expressing LifeAct-mCherry to visualize filamentous actin (Fig. 4A and Movies S1, 2). Similar to what we observed in *Tubb6* KO osteoclasts (Maurin et al., 2021), the actin ring in *Arhgap10* KO osteoclasts appeared unstable (Movies S1, 2). To quantify the effect of *Arhgap10* KO on actin ring stability, we compared the position of LifeAct-mCherry labelling between time points separated by 2 min. The median overlap frequency of LifeAct-mCherry staining between time points was 0.61 in control osteoclasts, whereas this value dropped significantly to 0.46 and 0.49 respectively in *Arhgap10* KO osteoclasts from clones #24 and #28 (Fig. 4B) showing that the KO of *Arhgap10* destabilizes the actin ring. These data demonstrate that ARHGAP10 regulates the stability of the actin ring in osteoclasts.

**Figure 4:**
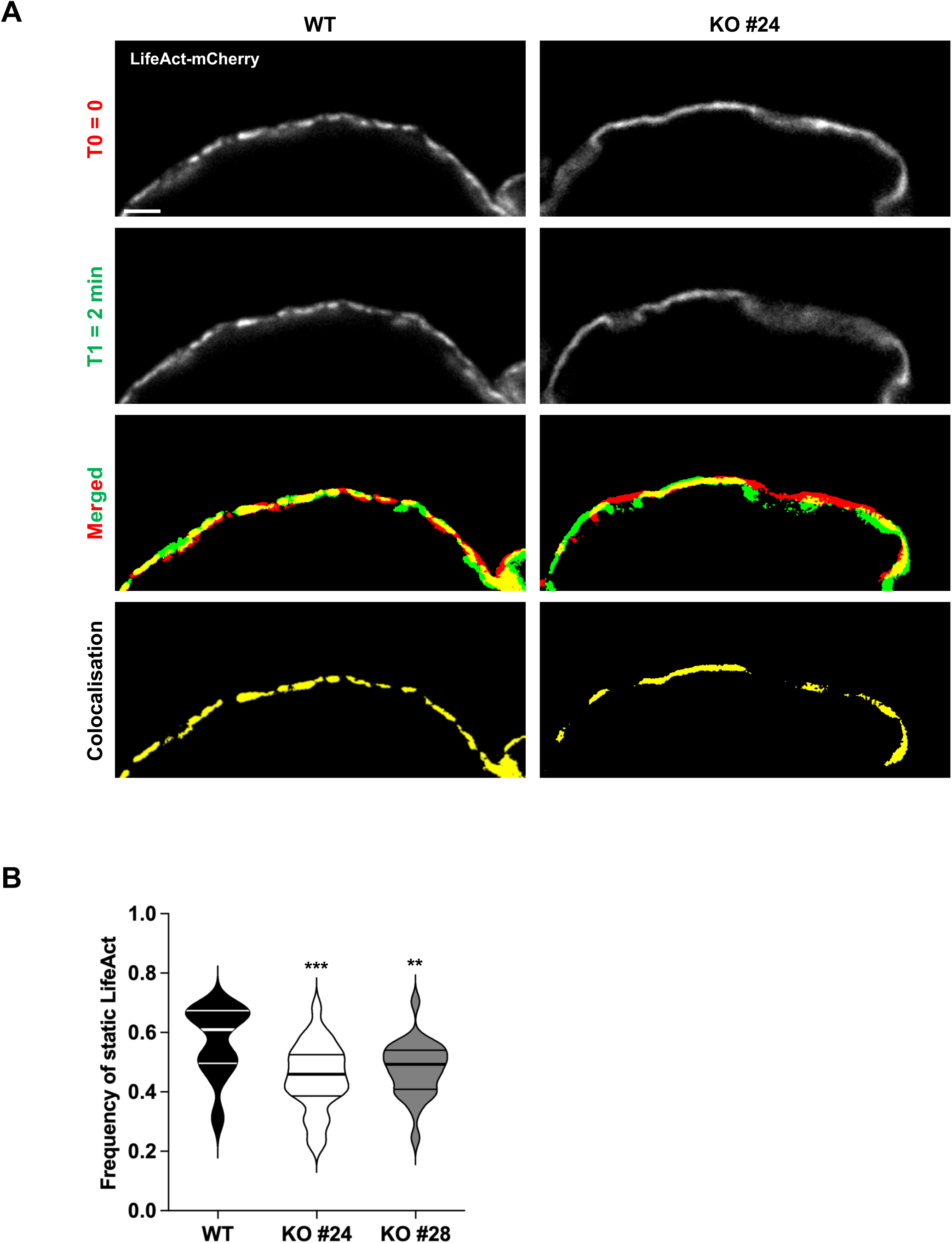
Arhgap10 KO affects actin ring dynamics. **(A)** Representative live images of osteoclasts derived from WT or *Arhgap10* KO cells expressing LifeAct-mCherry grown on plastic dishes. The localization of LifeAct-mCherry signal at T0 = 0 min (red) and T1 = 2 min (green), with the overlapping areas in yellow on the merged images and isolated on the colocalization images. Bar, 20 µm. **(B)** Violin plot representing the frequency of static LifeAct-mCherry that persists at the same position during two minutes timelapses. 22 WT, 27 Arhgap10 #24 KO and 25 Arhgap10 #28 KO osteoclasts from three independent experiments were analyzed. One-way Anova test: ***p < 0.001 and **p < 0.01 (see Experimental Procedures for calculation details).

### The microtubule binding-capacity of ARHGAP10 participates in osteoclast activity

Our data demonstrated that ARHGAP10 is a new MAP important for osteoclast activity by regulating actin ring organization and dynamics. In order to evaluate the implication of microtubule-binding capacity of ARHGAP10 in osteoclast activity, we performed complementation experiments to re-express full-length myc-tagged ARHGAP10, either WT or K37,41,44E, in *Arhgap10* KO osteoclasts. For this, the tetracycline inducible constructs were stably introduced in RAW264.7 *Arhgap10* KO #28 cells using the *Sleeping Beauty* transposon system (Mátés et al., 2009). Addition of doxycycline induced the expression of exogenous myc-tagged ARHGAP10 proteins in *Arhgap10* KO osteoclasts, nevertheless at higher levels than endogenous ARHGAP10 in WT osteoclasts (Fig. 5A). In order to confirm that exogenous ARHGAP10 proteins behave as expected in terms of microtubule binding, tubulin/microtubule fractionation of osteoclasts was performed. In agreement with the results obtained in HEK293T cells (Fig. 2D), around 50% of ectopic WT ARHGAP10 associated with the microtubule containing fraction while the K37,41,44E mutant was barely detectable in this fraction (Fig. S2). This result confirms that the K37,41,44E mutation in the BAR domain affects ARHGAP10 binding to microtubules in osteoclasts. We then compared the ability of *Arhgap10* KO osteoclasts, expressing or not the ectopic myc-tagged ARHGAP10, to dissolve hydroxyapatite. We found that upon complementation with WT ARHGAP10, the activity of *Arhgap10* KO osteoclasts was stimulated by 5.9-fold, whereas it was only increased by 1.6-fold when complemented with ARHGAP10 K37,41,44E mutant that cannot bind to microtubules (Fig. 5B). We tested whether ARHGAP10 overexpression could also rescue the activity of *Tubb6* KO osteoclasts, which is inhibited by around 70 % compared to WT osteoclasts (Maurin et al., 2021). For this, we expressed myc-tagged ARHGAP10 constructs in RAW264.7 *Tubb6* KO osteoclasts. The overexpression of exogenous the ARHGAP10 proteins in *Tubb6* KO osteoclasts was efficient, as assessed by immunoblot analysis (Fig. 6A). However, neither the overexpression of ARHGAP10 WT or K37,41,44E modified the resorption activity of *Tubb6* KO osteoclasts (Fig. 6B). Thus, ARHGAP10 overexpression does not rescue the absence of TUBB6 in osteoclasts.

**Figure 5:**
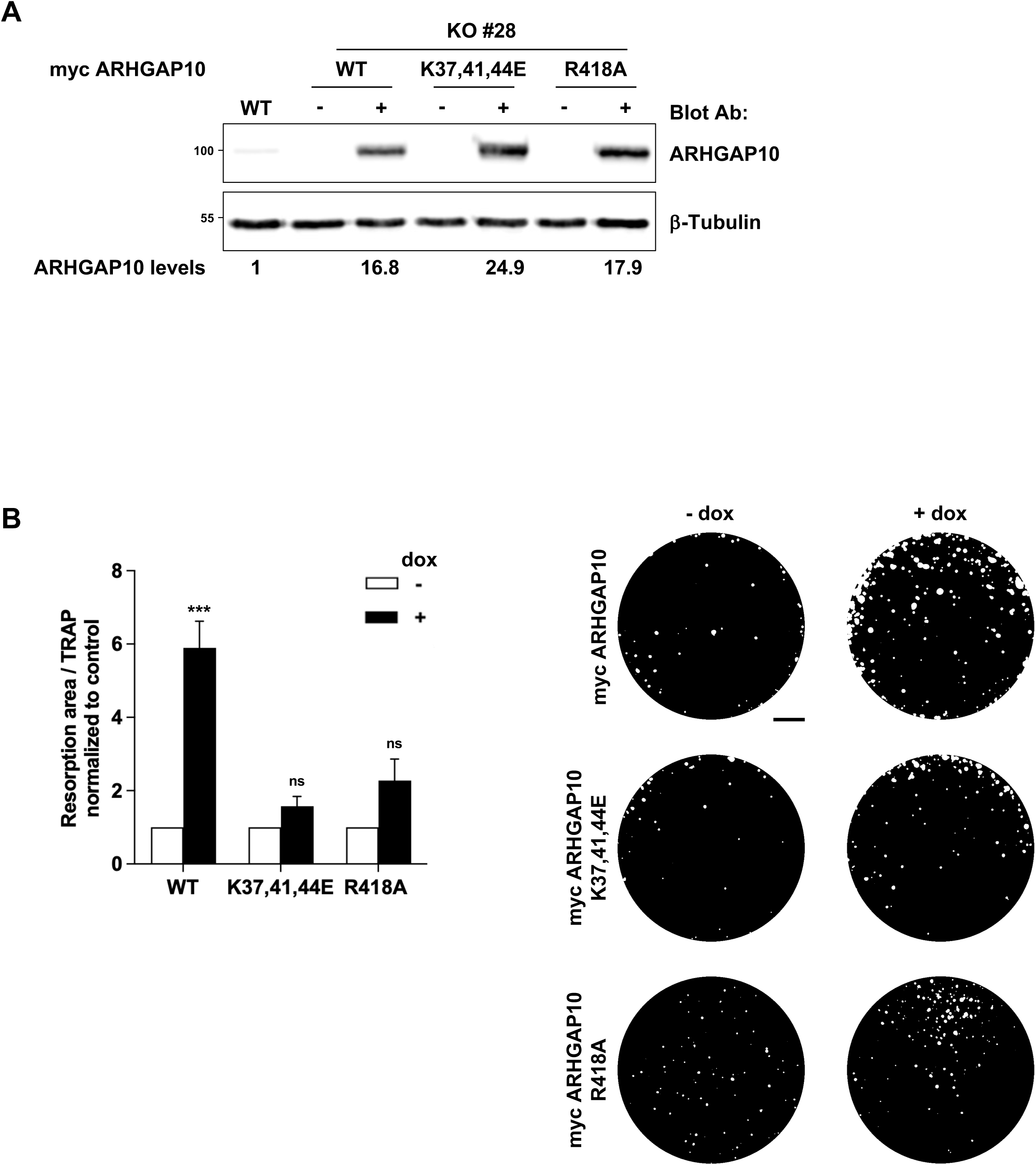
Microtubule binding and GAP activity of ARHGAP10 are both required for its activity in osteoclasts. **(A)** Representative immunoblot analysis of lysates from WT or *Arhgap10* #28 KO osteoclasts expressing (+) or not (-) myc-ARHGAP10 WT, K37,41,44E or R418A. β-tubulin shows equal protein loading. **(B)** Graph representing the average specific resorption activity of *Arhgap10* KO osteoclasts expressing (black bars, + dox) the indicated ARHGAP10 constructs, normalized to the activity of the corresponding non-expressing (white bars, - dox) osteoclasts (n = 12 for ARHGAP10 WT and K37,41,44E and n=4 for R418A, ± SEM; 8 wells analyzed per experiments). Wilcoxon test: ***p < 0.001, not significant (ns). Representative images of von Kossa staining of the resorption wells are presented on the right. Bar, 1 mm.

**Figure 6:**
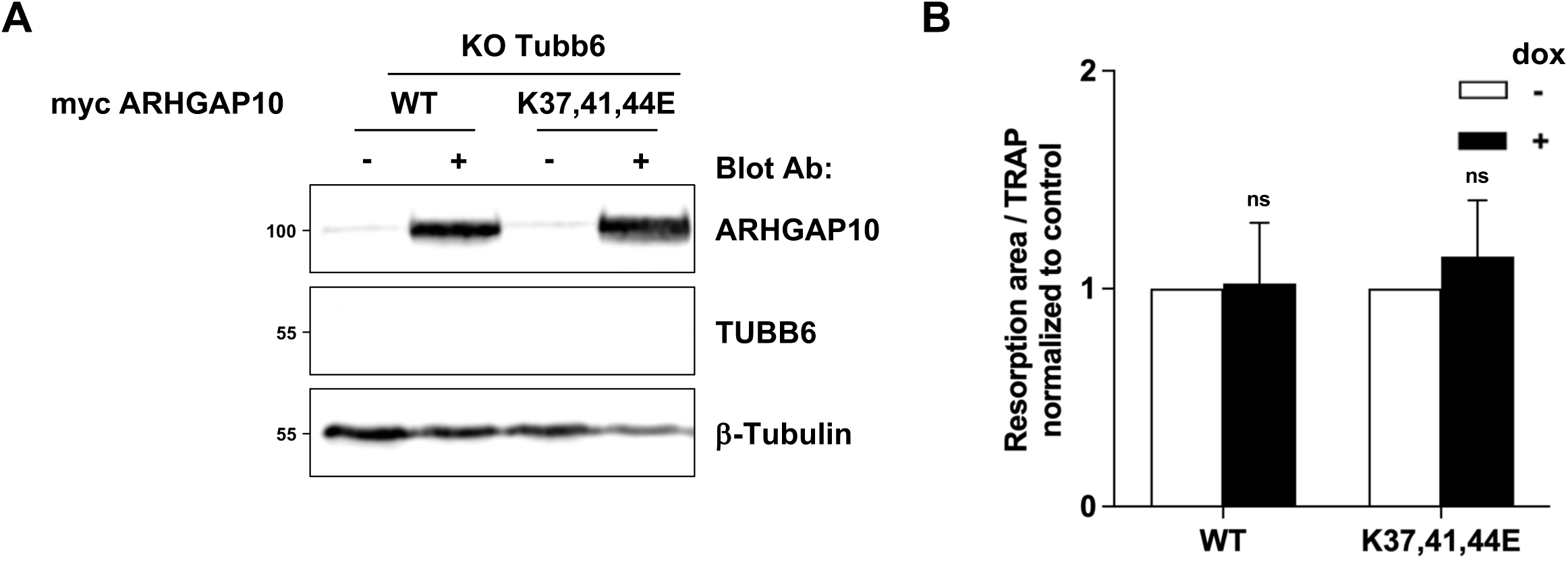
ARHGAP10 does not complement *Tubb6* KO in osteoclasts. **(A)** Representative immunoblot analysis of lysates from WT or *Tubb6* KO osteoclasts expressing (+) or not (-) myc-ARHGAP10 WT or K37,41,44E. β-tubulin shows equal protein loading. **(B)** Graph representing the average specific resorption activity of *Tubb6* KO osteoclasts expressing (black bars, + dox) the indicated ARHGAP10 constructs, normalized to the activity of the corresponding non-expressing (white bars, - dox) osteoclasts (n = 6, ± SEM; 8 wells analyzed per experiments). Wilcoxon test: not significant (ns).

ARHGAP10 is a GAP for CDC42 and RHOA both being essential for osteoclast activity. Thus, we investigated whether the GAP activity of ARHGAP10 is important for osteoclast function. We introduced a tetracycline inducible construct expressing a GAP-dead mutant of ARHGAP10 (R418A) (Li et al., 1997) in RAW264.7 *Arhgap10* KO #28 cells, as described above. Myc-tagged ARHGAP10 R418A protein was specifically detected upon doxycycline treatment (Fig. 5A). The behavior of this mutant in terms of microtubule-binding was assessed by tubulin/microtubule fractionation of osteoclasts. The R418A mutation did not alter the association of ARHGAP10 with the microtubule fraction (Fig. S2), in agreement with the above finding that the GAP-SH3 fragment of ARHGAP10 is not able to bind microtubles (Fig. 1C). We then assessed the ability of ARHGAP10 R418A to complement the hydroxyapatite dissolution defect of *Arhgap10* KO osteoclasts. The induction of ARHGAP10 R418A expression only increased by 2.5-fold the activity of osteoclast, as compared to the 5.9-fold increase upon expression of the WT protein (Fig. 5B).

Altogether, these results suggest that the microtubule-binding capacity and the GAP activity of ARHGAP10 are both necessary for the activity of osteoclasts.

## DISCUSSION

Actin cytoskeleton and microtubules cooperate for efficient bone resorption by osteoclasts (Destaing et al., 2003; Morel et al., 2024). We recently demonstrated that the isotype 6 of tubulin-β (TUBB6) is important for actin and microtubule crosstalk in osteoclasts and identified ARHGAP10 as a protein whose binding to microtubules may depend on TUBB6 (Guérit et al., 2020; Maurin et al., 2021). In the present study, we showed that ARHGAP10 is a *bona fide* MAP, which binds microtubules through its BAR-PH domain. Additionally, we show that ARHGAP10 is essential for osteoclast function, which relies on both ARHGAP10 ability to bind microtubules and its GAP activity.

ARHGAP10 is a negative regulator of CDC42 and RHOA GTPases, which are both involved in actin ring formation and dynamics in osteoclasts. Indeed, CDC42 is crucial for actin polymerization required for podosome formation and thus for actin ring (Ito et al., 2010). On the other hand, fine tuning of RHOA activity is necessary for correct organization of osteoclast actin ring (Destaing et al., 2003; Morel et al., 2024). We show here that in *Arhgap10* KO osteoclasts, the dynamics of the actin ring is affected and actin rings are smaller, which is in line with former results obtained using siRNAs (Steenblock et al., 2014). We further found that the lack of ARHGAP10 severely reduces the activity of osteoclasts. Thus, this negative regulator of CDC42 and RHOA is essential for osteoclast activity. Eventhough the GAP activity of ARHGAP10 was required for efficient rescue of resorption-activity of *Arhgap10* KO osteoclasts, we could not detect any change in the total activity of CDC42 and RHOA upon the knock-out of *Arhgap10* KO, assessed by pull down assays (Fig. S3). As neither *Arhgap10* (this study) nor *Tubb6* KO (Maurin et al., 2021) provoked a change in the total activity of CDC42 and RHOA, one should consider a subtler local regulatory effect on the activity of the GTPases. It would be of interest to study the local regulatory effect of ARHGAP10 using biosensors for the activity of CDC42 and RHOA for instance using FRET microscopy, a technic never used in osteoclasts so far.This suggests that the regulatory role of ARHGAP10 on CDC42 and/or RHOA activities in osteoclasts could be local.

We initially identified ARHGAP10 as an potential effector of TUBB6 activity in osteoclasts, as we had found that more ARHGAP10 appeared to associate with microtubules in the absence of TUBB6 (Maurin et al., 2021). Here we show that ARHGAP10 depletion induces similar actin ring and resorption defects as in *Tubb6* KO osteoclast, suggesting that increasing the binding of ARHGAP10 to microtubules has similar consequences to depleting ARHGAP10. Overexpression of TUBB6 has a microtubule-destabilizing effect (Bhattacharya and Cabral, 2004, 2009; Maurin et al., 2021); conversely, we reported that *Tubb6* KO in osteoclasts decreases microtubule dynamics (Maurin et al., 2021). Thus, microtubule destabilization by TUBB6 may regulate ARHGAP10 activity by favouring its local release from microtubules. In line with this hypothesis, the ectopic expression of ARHGAP10 cannot complement resorption defects of *Tubb6* KO osteoclasts. In agreement with the important regulatory role of microtubules in ARHGAP10 function is that the microtubule-binding defective K37,41,44E mutant of ARHGAP10 is less efficient than the WT at rescuing the resorption activity of *Arhgap10* KO osteoclasts. Therefore, ARHGAP10 binding to microtubules regulated by TUBB6 could locally modulate the activity of CDC42 and/or RHOA for proper osteoclast activity.

We show here that ARHGAP10 is a bona fide MAP, which binds microtubules via its BAR-PH domain. The binding of MAPs to microtubules is usually electrostatic and involves positively charged residues that interact with the intrinsically disordered and negatively-charged C-terminal tails of α- and β-tubulins at the surface of the microtubule lattice (Roll-Mecak, 2020; Bigman and Levy, 2020). By modelling the structure of the BAR-PH domain of ARHGAP10, we could identify several exposed positively charged residues enriched within the concave side of the BAR domain. As the mutation of three lysine residues to negatively charged residues (K37,41,44E) almost completely abolished microtubule binding, it is very likely that the concave face of ARHGAP10 BAR domain can interact with negatively-charged C-terminal tails of α- and β-tubulins on microtubules. The concave side of BAR domains is known to sense membrane curvature and to induce membrane tubulation (Salzer et al., 2017) but it also represents a binding platform for a variety of proteins (Carman and Dominguez, 2018). A BAR domain can also accommodate a polymer such as a microtubule: indeed, CDC42 interacting protein 4 (CIP4), a CDC42-effector, has been shown to bind microtubules *in vitro* through its amino-terminal F-BAR domain (Tian et al., 2000). The F-BAR domain of CIP4 is fairly flat (Shimada et al., 2007) as compared to the structural model of the BAR-PH domain of ARHGAP10. Still, the BAR domain of ARHGAP26, which is 67.8% identical with that of ARHGAP10 (Fig. S1D), has been shown to bind small 50 nm-diameter liposomes (Lundmark et al., 2008). As the diameter of a microtubule is only 25 nm (Roll-Mecak, 2020), the narrow angle of the BAR-PH domain of ARHGAP10 from the proposed structural model should easily accommodate a microtubule, all the more as it interacts with the intrinsically disordered tails of α- and β-tubulins that stick out of the microtubule.

BAR domains are sensing membrane curvature and some are inducing membrane tubulation. Membrane binding by BAR domains involves electrostatic interactions and positively charged residues located within the concave face. The crystal structure of the BAR domain of Amphiphysin and mutation analysis highlighted the crucial role of positively charged residues located within helix 2 in membrane binding (Peter et al., 2004). In the case of Centaurin β2, which contains a PH domain downstream of the BAR domain, similar to ARHGAP10, the membrane binding is mostly mediated by the PH domain but sensing membrane curvature and tubulation relies on the BAR domain and, in particular, on two lysine residues (K124 and 125) located within helix 2 (Peter et al., 2004). Mutating corresponding lysine residues (K131 and 132) in the BAR domain of ARHGAP26 prevents membrane tubulation *in vitro* and tubule association in HeLa cells of the BAR-PH (Lundmark et al., 2008). The BAR-PH domain of ARHGAP10 has recently been shown to bind and to induce tubular endosomes, as part of a MICAL1-ARHGAP10-WDR44 complex, in HeLa cells (Lucken-Ardjomande Häsler et al., 2020). Similar to ARHGAP26, this activity of ARHGAP10 likely depends on lysine residues 131 and 132 of the BAR domain (indicated by purple dots in Fig. S1D). Lysine residues involved in microtubule binding (K37,41 and 44 indicated by blue dots in Fig. S1D) are all located within helix 1 of the BAR domain and their mutations to glutamate residues are therefore unlikely affecting membrane tubulation and tubule binding but it deserves to be further investigated.

Altogether our data demonstrate that ARHGAP10 is a novel MAP regulating osteoclast activity. Interfering with ARHGAP10 function, for instance by targeting the ARHGAP10-microtubule interface and/or GAP activity, could represent a therapeutic strategies against pathological bone loss.

## EXPERIMENTAL PROCEDURES

### Cell lines and lentivirus production

RAW264.7 cells were a gift from Kevin P McHugh (Gainesville, FL, USA). Human embryonic kidney (HEK293T) and HeLa cells were obtained from the ATCC. All cell lines were maintained in DMEM (Life Technologies) supplemented with 10% heat inactivated fetal calf serum (Eurobio), Penicillin-Streptomycin (Life Technologies), 2 mM glutamine (Life Technologies). Lentiviral particles were produced at the Plateforme de Vectorologie de Montpellier (https://www.pvm.cnrs.fr/, Montpellier, France).

### DNA constructs

Vectors encoding myc full-length human ARHGAP10 was generated by cloning the corresponding synthetic cDNA, ARHGAP10 WT or K37,41,44E (synthetized by GenScript Biotech, Netherlands), in pRK5-myc. R418A mutation was introduced in pRK5-myc ARHGAP10 WT using QuickChange Site-Directed Mutagenesis Kit (Agilent Technology). Vectors encoding HA GST or myc BAR-PH (aa, 1-388) and GAP-SH3 (aa, 373-786) were generated by PCR-based cloning in pRK5-HA GST or pRK5-myc using synthetic ARHGAP10 WT or K37,41,44E as templates. Corresponding cDNA were cloned into pET28a to express His GST BAR-PH or His GST GAP-SH3 in bacteria. Vector encoding inducible myc ARHGAP10 WT, K37,41,44E or R418A were generated by PCR-based cloning in pSBtet-Bla (a gift from Eric Kowarz, Addgene plasmid #60510; http://n2t.net/addgene:60510; RRID:Addgene_60510) using pRK5-myc ARHGAP10 WT or K37,41,44E as templates (Kowarz et al., 2015). All constructs were sequenced.

### Osteoclast differentiation and activity

For osteoclast differentiation, RAW264.7 cells between passages 2 and 12 were cultured in a humidified incubator at 37 °C and 5% CO_2_, in α-MEM supplemented with 10% heat inactivated foetal calf serum (Biowest), Penicillin-Streptomycin (Life Technologies), 2 mM glutamine (Life Technologies) and 50 ng/mL RANKL (Peprotech). Medium was changed every two days until osteoclast differentiation at day 4 or 5. After 3 days of differentiation, mononucleated cells were removed by washes with PBS and osteoclasts were detached by treatment with Accutase™ (Sigma Aldrich) followed by scrapping. Detached osteoclasts were transferred on Apatite Collagen Complex (ACC)-coated coverslips, prepared as reported previously (Maurin et al., 2018) and analyzed 48 hours later.

To measure their activity, osteoclasts were detached from the culture plate as described above and seeded for 24 hours onto uncoated 96-well plates or for 48 hours onto inorganic crystalline calcium phosphate (CaP)-coated 96-well plates prepared as described (Cen et al., 2022). For each condition, 8 wells of uncoated 96-well plates were stained for tartrate resistant acid phosphatase (TRAP) activity to measure osteoclast surface and 8 wells in (CaP)-coated 96-well plates were stained with von Kossa to measure CaP dissolution as described previously (Brazier et al., 2009). When appropriate, TRAP activity was also measured in the supernatant of cells growing onto (CaP)-coated 96-well plates as described (Remmers et al., 2023) with minor modifications. Briefly, 100 µl of cell supernatant was mixed with 100 µl of TRAP reaction buffer (100 mM sodium acetate, 50 mM sodium tartrate, pH 5.5 containing 5 mg/ml 4-nitrophenylphosphate) and incubated at 37°C for 60 minutes. Reaction was stopped by addition of 50 µl 0.5 M NaOH and absorbance was measured at 405 nm (specific) and 490 nm (unspecific) using a POLARstar Omega plate reader (BMG Labtech). TRAP activity was defined as OD_405_-OD_490_. TRAP and von Kossa-stained wells were imaged with a Leica Z16 APO macroscope equipped with CoolSNAP™ fx CCD camera (Photometrics) and quantifications were carried out with ImageJ2 (2.16.0/1.54p) software. In each experiment, osteoclast specific activity was expressed as the average area resorbed in 8 wells stained with von Kossa divided by the average osteoclast surface in 8 wells stained with TRAP or by the average TRAP activity measured in the supernatant of 8 wells; this value was then normalized to the specific activity of the control condition of the experiment.

### Generation of Arhgap10 knock-out RAW264.7 cells and variants

A guide RNA (gRNA) targeting the eighth exon of mouse *Arhgap10* gene (5’-AGAAGGCTACCTGTACGTCC-3’) was designed and cloned in lentiCRISPRv2 vector, which also provides the expression of the *Streptococcus pyogenes* Cas9 and puromycin resistance a gift from Feng Zhang (Addgene plasmid # 52961; http://n2t.net/addgene:52961; RRID:Addgene_52961) (Sanjana et al., 2014). Control empty and *Arhgap10* gRNA-containing lentiCRISPRv2 were used to produce lentiviral particles and to generate respectively control (WT) and *Arhgap10* KO RAW264.7 cells, as previously described for *Tubb6* (Maurin et al., 2021). ARHGAP10 expression was monitored by immunoblot analysis using home-made polyclonal antibodies. Rabbit polyclonal antibodies against GST-tagged SH3 from human ARHGAP10 (718-786) were raised at the CRBM animal facility and affinity purified as described (Lucken-Ardjomande Häsler et al., 2014). Selected clones of RAW264.7 cells were then used to differentiate WT and *Arhgap10* KO osteoclasts.

RAW264.7 cells KO for *Arhgap10* (clone #28) stably expressing myc ARHGAP10 WT, K37,41,44E or R418A under a doxycycline-inducible promoter were generated using the *Sleeping Beauty* transposon system by co-transfecting pSBtet-Bla-myc ARHGAP10 WT, K37,41,44E or R418A and pCMV(CAT)T7-SB100 (a gift from Zsuzsanna Izsvak, Addgene plasmid # 34879; http://n2t.net/addgene:34879; RRID:Addgene_34879) (Mátés et al., 2009), at a 5/1 ratio using jetPEI^®^ following manufacturer instructions (Polyplus transfection). 48 hours after transfection cells were selected with 10 µg/ml blasticidin and later maintained as a pool. Protein expression was induced with 100 ng/ml doxycycline for 48 hours after three days of differentiation.

### Immunoblot analysis

Osteoclast lysates were either prepared in protein loading buffer (80 mM Tris-HCl; pH 6.8, 90 mM DTT, 2% SDS, 10% glycerol, 0.004 % bromophenol blue and 10 mM iodoactetamide) or in lysis buffer (10 mM NaH_2_PO_4_, 100 mM NaCl, 5 mM EDTA, 1% Triton X-100, 0.5% IGEPAL CA-630, 80 mM β-Glycerophosphate, 1 mM DTT, 50 mM NaF, 1 mM Na_3_VO_4_, and protease inhibitor cocktail). Protein concentrations were determined using Protein Assay Dye Reagent (BioRad) and BSA was used as standard. After resolution by SDS-PAGE, proteins were transferred to PVDF-FL membranes (Immobilon-FL Transfer Membrane) and analyzed by immunoblot. Primary antibodies were: ARHGAP10 (1:1000), Abcam antibodies Vinculin (ab108620, 1:2000) and GEF-H1 (ab155785, 1:500), Developmental Studies Hybridoma Bank antibodies total β-tubulin (E7, 1:5000) and total α-tubulin (12G10, 1:5000), Santa Cruz Biotechnologies antibodies total α-tubulin (YOL1/34, sc-5330, 1:2000), Cell Signaling Technology Myc-Tag (71D10, 2278, 1:2000), RhoA (67B9, 2117, 1:1000), BD Biosciences His (F24-796, 552565, 1:10000) and Abnova Cdc42 (M152, MAB1376, 1:500). Secondary antibodies were: Goat-anti-mouse Dylight800 (Invitrogen, SA535521), Goat-anti-rabbit Dylight800 (Invitrogen, SA535571), Horse-anti-mouse HRP-linked (Cell Signaling Technology, 7076), Goat anti-rabbit HRP linked (Cell Signaling Technology, 7074), Goat-anti-rat HRP-linked (Cell Signaling Technology, 7077). Signals were acquired and measured using the Odyssey Fc Imager (LI-COR Biosciences, USA).

### Immunofluorescence and microscopy

All imaging was performed at Montpellier Ressources Imagerie (MRI) in Montpellier (https://www.mri.cnrs.fr). Osteoclasts grown for 48 hours on ACC were fixed for 20 minutes at 37°C in 4% paraformaldehyde and 10 µM Paclitaxel (Sigma Aldrich) in PHEM (60 mM PIPES, 25 mM HEPES, 10 mM EGTA, 2 mM MgCl_2_, pH 6.8) and cells were then permeabilized with 0.1% Triton X-100 in PBS for 4 minutes. After blocking for 30 minutes with 1% BSA in PBS, cells were processed for F-actin and DNA staining using respectively Phalloidin rhodamine (Sigma Aldrich, P1951, 1:10,000) and Hoechst 33342 (Sigma Aldrich, 14533, 1:10,000). Coverslips were mounted in Citifluor MWL4-88-25 (CliniSciences) and imaged with Zeiss Axioimager Z2 widefield microscope equipped with oil objective 63× Plan Apochromat 1.4 NA oil, Orca Flash 4.0 V2+ (Hamamatsu) camera driven by Zen 2 (Zeiss).

### Image quantification and live imaging

Actin organization was evaluated visually after F-actin staining in fixed osteoclasts seeded on ACC: an actin ring was classified as abnormal when F-actin staining was fragmented, weak, or absent in more than half of the periphery of the sealing zone, or multiple when more than 4 small sealing zones per osteoclast were present.

For live imaging, osteoclasts were differentiated from WT and *Arhgap10* KO RAW264.7 cells expressing LifeAct-mCherry to follow actin dynamics as described (Maurin et al., 2021). LifeAct-mCherry-expressing osteoclasts were imaged at 37°C and 5% CO_2_ with Olympus IX83 microscope equipped with scMOS ZYLA 4.2 MP camera, one image every 2 minutes, with air objective 20× LUCPLFLN 0.45 NA PH1 driven by Metamorph. The overlap of actin ring position between different time points was determined using ImageJ2 (2.16.0/1.54p) as described (Maurin et al., 2021). A minimum of 5 cells per condition were considered and a minimum of 10 measurements per cell were done and averaged. The graphical representation and statistic analysis were performed using those averaged values obtained from three independent experiments.

### Microtubule binding

Microtubule co-sedimentation assays using cell extracts derived from osteoclasts were performed as described (Maurin et al., 2021). Briefly, osteoclasts were lysed on ice in 1X BRB80 (80 mM PIPES, 1 mM EGTA, 1 mM MgCl_2_, pH 6.8) supplemented with 1 mM DTT, protease inhibitor cocktail, and 0.1% IGEPAL CA-630. After clarification at 100,000 × g, 100 µg of lysate supplemented with 2 mM GTP was incubated 5 min on ice followed by 10 min at 37°C, and then 10 minutes more after addition of 20 mM paclitaxel. After centrifugation at 100,000 × g over a 40% glycerol cushion in 1X BRB80 supplemented with 20 mM paclitaxel, the supernatant was collected and, after rinsing, the pellet was resuspended in the same volume as the supernatant. Protein association with microtubules was also assessed by cell fractionation into tubulin and microtubule fractions as described (Bompard et al., 2018) from HEK293T expressing various ARHGAP10 constructs 24 hours after transfection using jetPEI^®^ following manufacturer instructions (Polyplus transfection).

For *in vitro* microtubule co-sedimentation assay, His- and GST-tagged ARHGAP10 BAR-PH WT or K37,41,44E and GAP-SH3 domains were expressed in BL21 Rosetta2 pLysS bacteria and proteins were purified on HIS-Select^®^ Nickel Affinity Gel (Sigma-Aldrich) as previously described (Bompard et al., 2005). Eluted proteins were dialyzed against 50 mM Tris-HCl pH8, 100 mM NaCl, 1 mM DTT and concentrated using Amicon^®^ Ultra centrifugal filters (Millipore). 2 µM recombinant proteins were incubated with increasing concentrations of microtubules, polymerized from pig brain tubulin purified as described (Castoldi and Popov, 2003), in 1X BRB80 (80 mM PIPES, 1 mM EGTA, 1 mM MgCl_2_, pH 6.8) containing 1 mM DTT and 20 µM paclitaxel for 5 minutes at room temperature followed by a centrifugation through a cushion of 40% glycerol in 1X BRB80 containing 1 mM DTT and 20 µM paclitaxel at 100,000 × g for 15 minutes at 30°C. Supernatants were retrieved and pellets washed once with 1X BRB80 containing 20 µM paclitaxel before being resuspended in an equal volume of supernatant of loading buffer. The supernatant and pellet fractions were analyzed by immunoblot.

### CDC42 and RHOA pull-down assays

The activity of CDC42 and RHOA GTPases in osteoclasts was assessed by pull-down assays. The GTPase-binding domain (GBD, aa 124-274) of Bos Taurus N-WASP fused to GST was used to immobilize active CDC42 by pull-down assay as described (Bompard et al., 2005). The RHOA-binding domain (RBD) of Rhotekin fused to GST was used to immobilize active RHOA by pull-down assay as described (Morel et al., 2024).

### Statistical analyses

Statistical significance was assessed either with parametric statistical test after normality assessment or with non-parametric tests, with GraphPad Prism 10.4.1 (GraphPad Software, Inc.); p < 0.05 was considered statistically significant.

## DATA AVAILABILITY

All data are included within the article and supporting information.

## SUPPORTING INFORMATION

This article contains supporting information.

## ACKNOWLEDGMENTS

We acknowledge the imaging facility Montpellier Ressources Imagerie (MRI), a member of the national infrastructure France-BioImaging supported by the French National Research Agency (ANR-10-INBS-04, ‘Investments for the future’). We are very grateful to Juliette van Dijk and Andrey Kajava (CRBM Montpellier, France) for technical assistance, helpful advice and discussions. We want to especially thank Ewan McDonald (IRIM, Montpellier, France) for critical reading and editing of the manuscript. We also thank Drs Safa Lucken-Ardjomande Häsler and Harvey T. McMahon (Medical Research Council Laboratory of Molecular Biology, Cambridge, UK) for the generous gift of GST SH3-ARHGAP10 expression plasmid.

## AUTHOR CONTRIBUTIONS

Conceptualization: A.B., G.B; Methodology: A.B., G.B., L.J.; Validation: A.B., G.B., L.J.; Formal analysis: A.B., G.B; Investigation: A.B., G.B., L.J.; Writing - original draft: A.B., G.B.; Visualization: G.B.; Supervision: A.B., G.B.; Project administration: A.B., G.B.; Funding acquisition: A.B., G.B.

## FUNDING AND ADDITIONAL INFORMATION

This study was supported by the Centre National de la Recherche Scientifique (CNRS) and by the Université de Montpellier, and by grants from the GEFLUC (GEFLUC 276951 to G.B.), the Fondation ARC pour la Recherche sur le Cancer (PJA 20191209321 to A.B.) and the Agence Nationale de la Recherche (ANR-23-CE13-0033-01 to A.B.).

## LEGENDS OF SUPPORTING INFORMATION

**Figure S1:**
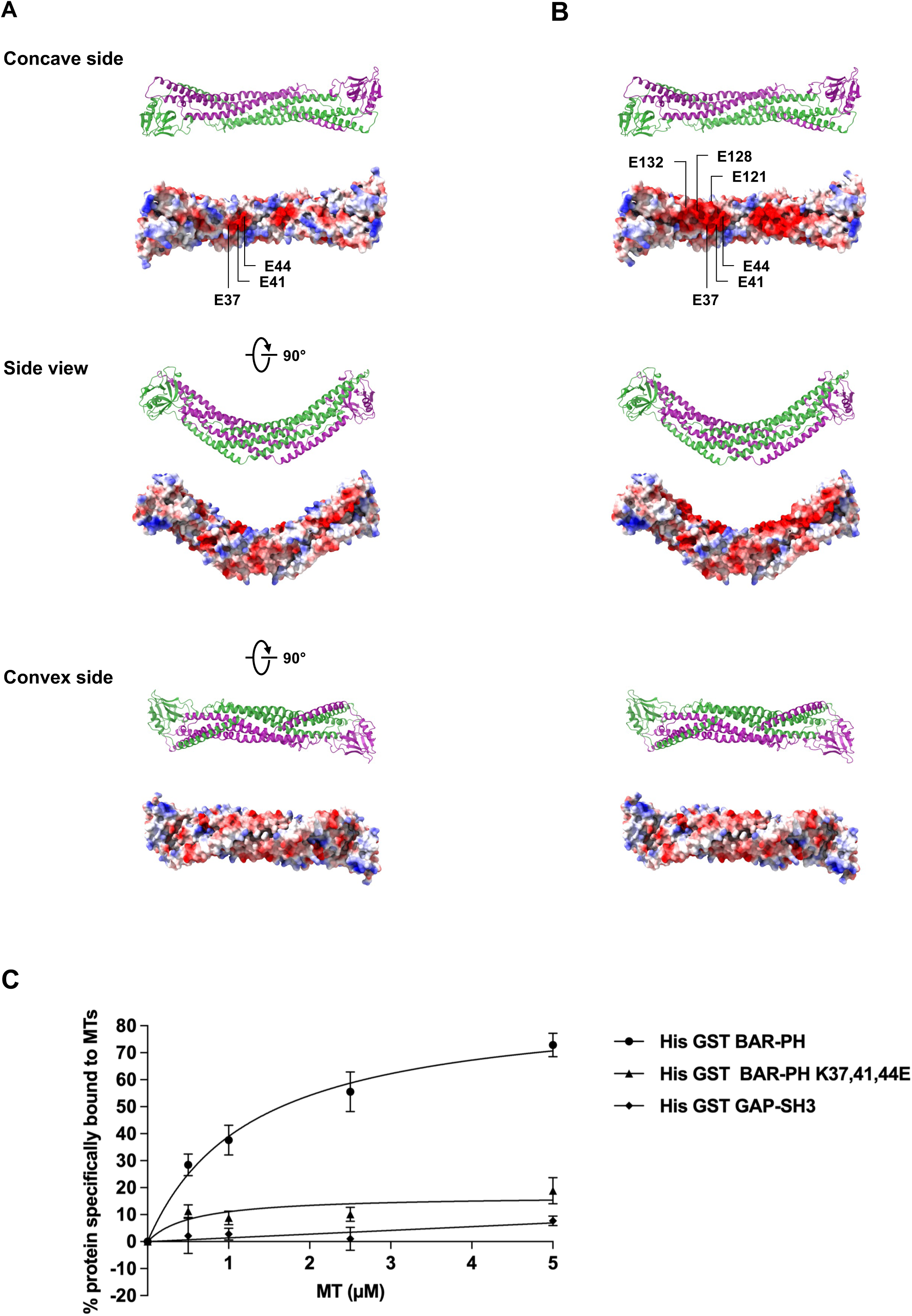
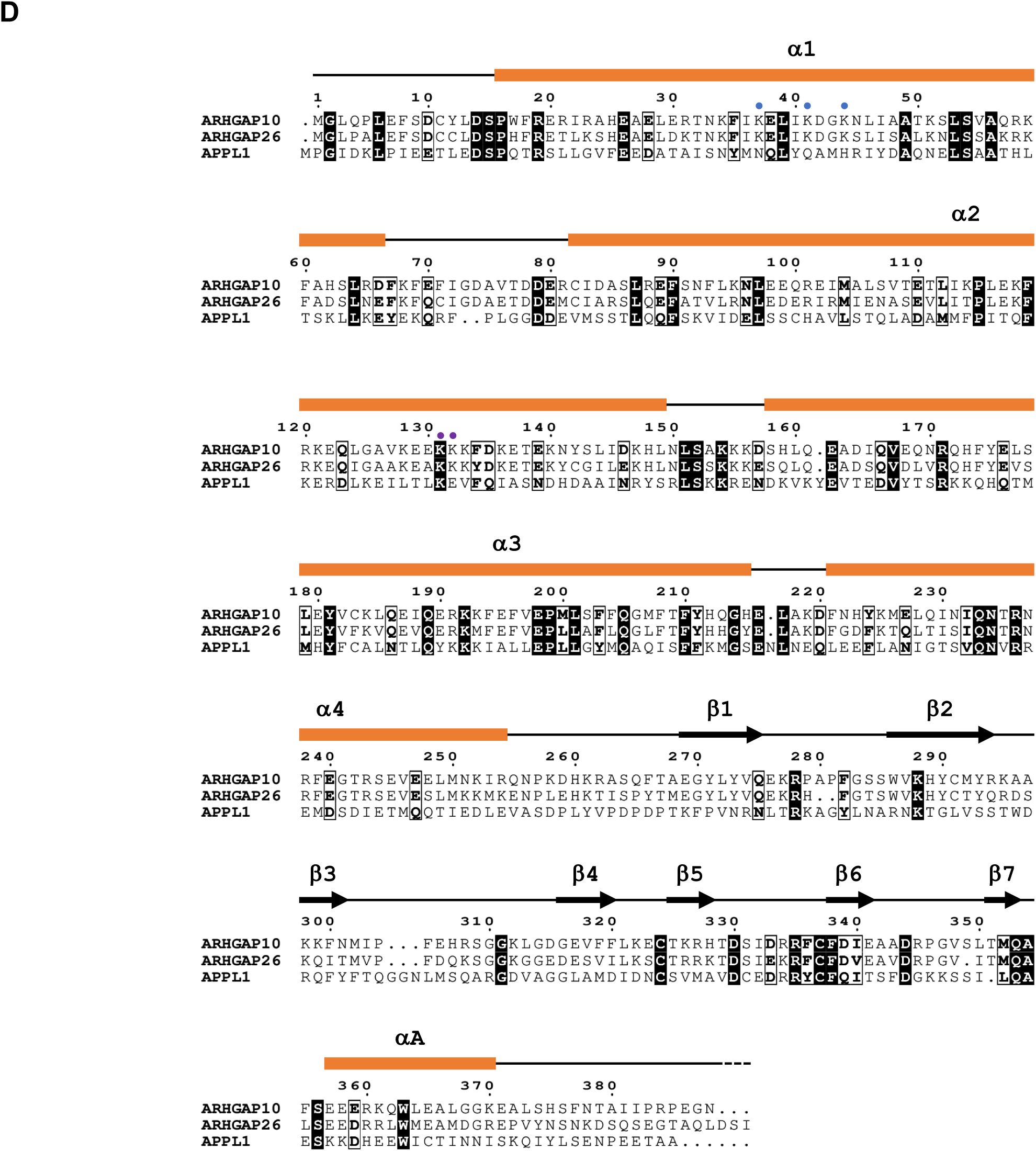
Effect of K37,41,44E mutation on structure and in vitro microtubule binding of ARHGAP10 BAR-PH domain. **(A)** Structural model of ARHGAP10 BAR-PH K37,41,44E mutant. The BAR-PH domain (1-388) of ARHGAP10 was modelized as in fig. 2A. The corresponding electrostatic surface of each side is represented bellow. Note that introduced mutations do not significantly affect the structure of the BAR-PH domain but the impact of indicated mutations on the charge is obvious mostly in the concave side. **(B)** Structural model of ARHGAP10 BAR-PH K37,41,44E and K121,128,132E mutant. This mutant was modelized and represented as in (A). **(C)** Graph quantifying the microtubule co-sedimentation of His GST BAR-PH K37,41,44E (n = 5, ± SEM) along with the curves corresponding to His GST BAR-PH and His GST GAP-SH3 presented in fig. 1D. **(D)** Sequence alignment of human ARHGAP10, ARHGAP26 and APPL1. The secondary structure elements are indicated. Conserved residues are boxed, identical residues are highlighted by black background and similar residues are bold. Blue and purple dots respectively show residues involved in microtubule binding and in sensing membrane curvature, inducing membrane tubulation, binding tubules.

**Figure S2:**
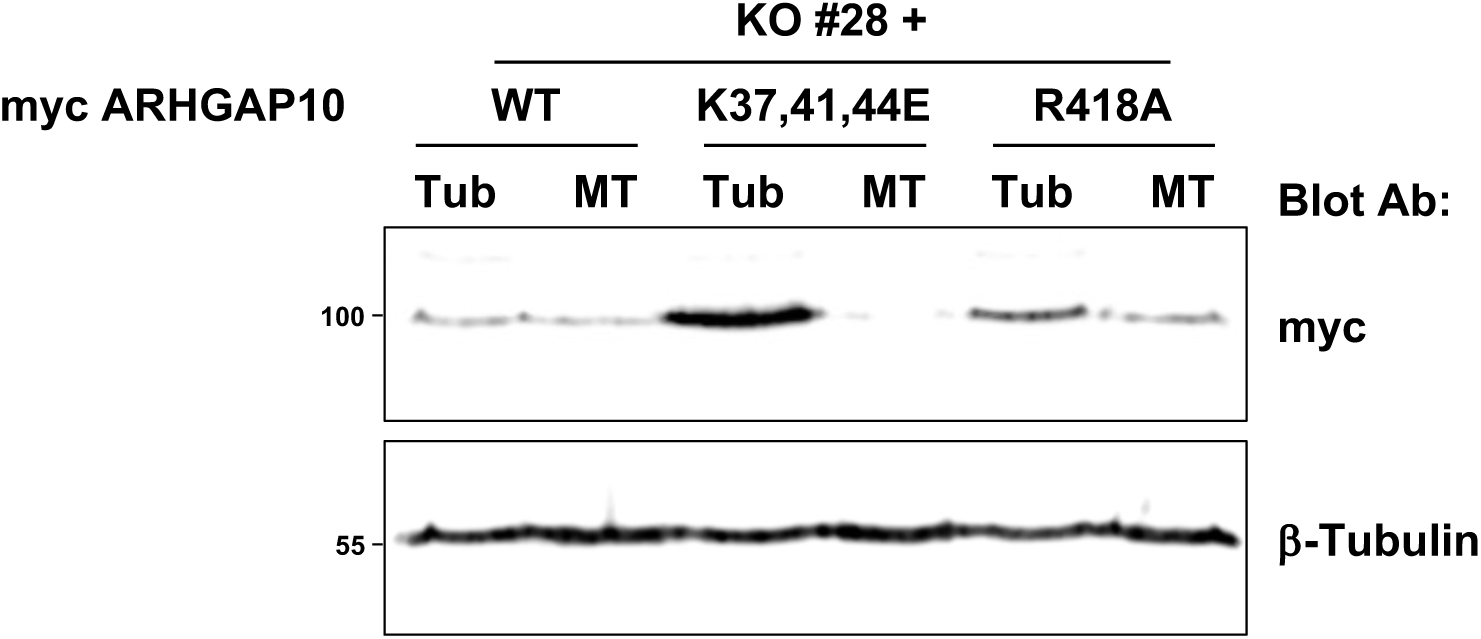
Microtubule association of exogenous ARHGAP10. Representative immunoblot analysis of lysates from Arhgap10 KO #28 osteoclast expressing myc-tagged ARHGAP10 WT, K37,41,44E or R418A subjected to tubulin/MT fractionation. Note that the association of ARHGAP10 with MT fraction is significantly inhibited by K37,41,44E mutations but not R418A mutation.

**Figure S3:**
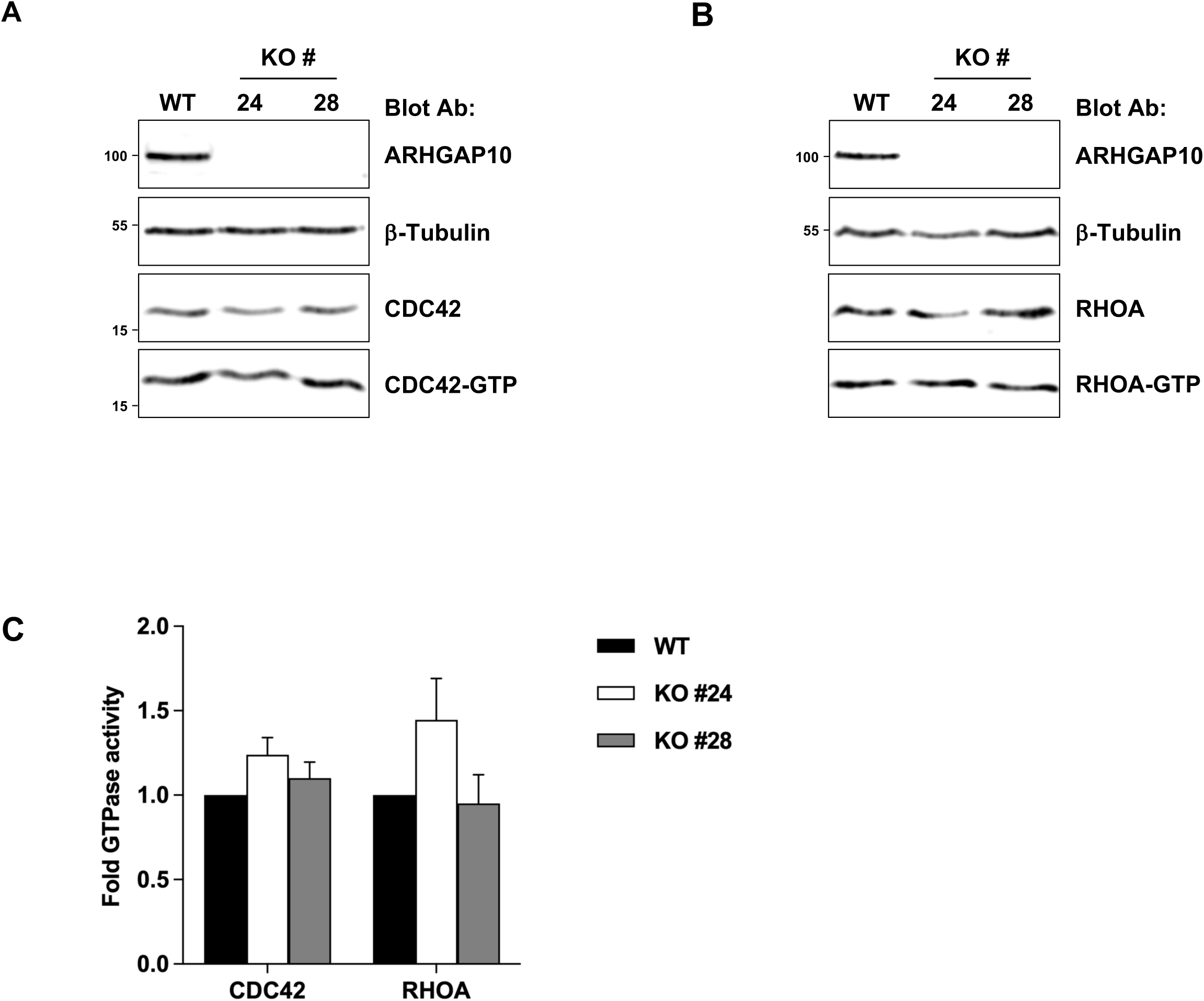
Global CDC42 and RHOA activities are not affected by ARHGAP10 depletion. **(A)** Representative immunoblot analysis of active CDC42 pull-down using the GTPase binding domain of N-WASP from WT and Arhgap10 KO osteoclasts. β-tubulin is used as a loading control and osteoclast differentiation is evaluated by mature CtsK. **(B)** Representative immunoblot analysis of active RHOA pull-down using the RHO binding domain of Rhotekin from same cells as in (A). **(C)** Graph representing fold changes in GTPase activity of osteoclast expressing (WT) or not ARHGAP10 (KO #24 or 28) normalized to the activity in control cells (WT) (n = 4 for CDC42 or 3 for RHOA, ± SEM).

**Movies S1 and S2.**

Live imaging of LifeAct-mCherry expressing WT (movie S1) or KO Arhgap10 #24 (movie S2) osteoclasts. Time is indicated as hour:min and scale bar is 20 µm.

